# An image-computable model of human visual shape similarity

**DOI:** 10.1101/2020.01.10.901876

**Authors:** Yaniv Morgenstern, Frieder Hartmann, Filipp Schmidt, Henning Tiedemann, Eugen Prokott, Guido Maiello, Roland W. Fleming

## Abstract

Shape is a defining feature of objects. Yet, no image-computable model accurately predicts how similar or different shapes appear to human observers. To address this, we developed a model (‘ShapeComp’), based on over 100 shape features (e.g., area, compactness, Fourier descriptors). When trained to capture the variance in a database of >25,000 animal silhouettes, ShapeComp predicts human shape similarity judgments almost perfectly (r^2^>0.99) without fitting any parameters to human data. To test the model, we created carefully selected arrays of complex novel shapes using a Generative Adversarial Network trained on the animal silhouettes, which we presented to observers in a wide range of tasks. Our findings show that human shape perception is inherently multidimensional and optimized for comparing natural shapes. ShapeComp outperforms conventional metrics, and can also be used to generate perceptually uniform stimulus sets, making it a powerful tool for investigating shape and object representations in the human brain.

## Introduction

One of the most important goals for biological and artificial vision is the estimation and representation of shape. Shape is the most important cue in object recognition [1–4] and is also crucial for many other tasks, including inferring an object’s material properties [5–9], causal history [10–13], or where and how to grasp it [14–18]. Shape is also central to many other disciplines, including computational morphology [19], anatomy [20], molecular biology [21], geology [22], meteorology [23], computer vision [24], and computer graphics [25]. It would be exceedingly useful to be able to characterize and quantify the visual similarity between different shapes (**Figure 1A**). Here we sought to build a model to estimate perceived shape similarity directly from images by integrating numerous shape metrics. Specifically, given a pair of shapes, {*f*_1_, *f*_2_}, the model should compare and combine shape metric *i* (of a total of *N*) to predict the perceived similarity between shapes, Ŝ, on a continuous scale (**Figure 1B**),

**Figure 1.**
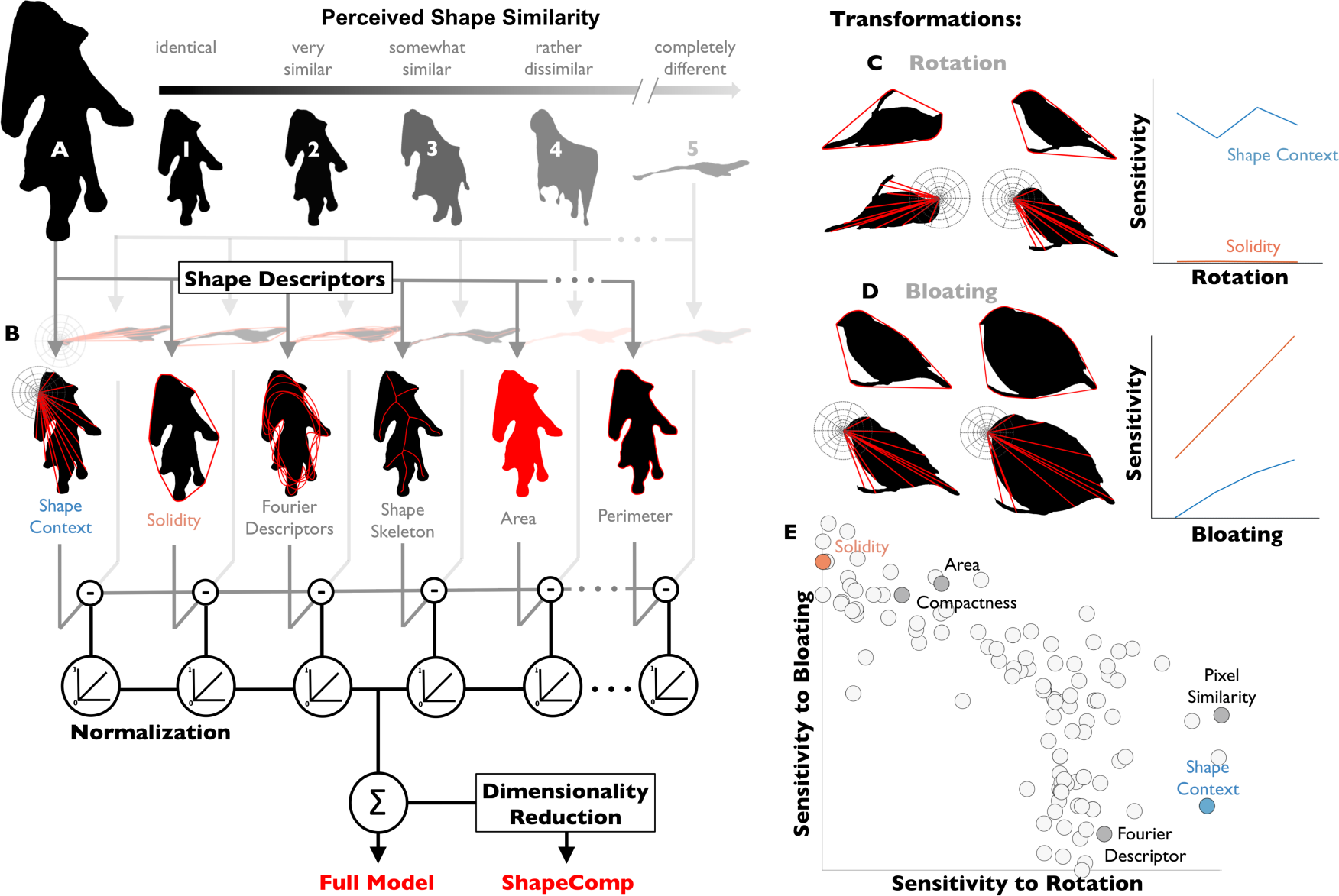
ShapeComp: a multidimensional perceptual shape similarity model. We readily perceive how similar shape (**A**) is from others (numbered 1-5). (**B**) Outline of our model, which compares shapes across >100 shape descriptors (6 examples depicted). The distance between shapes on each descriptor was scaled from 0 to 1 based on range of values in a database of 25,712 animal shapes. Scaled differences are then linearly combined to yield ‘Full Model’ response. Applying MDS to >330 million shape pairs from the Full Model yields a multidimensional shape space for shape comparison (‘ShapeComp’). We reasoned that many descriptors would yield a perceptually meaningful multidimensional shape space due to their complementary nature. (**C**) Some shape descriptors are highly sensitive to rotation (e.g., Shape Context), while (**D**) other descriptors are highly sensitive to bloating (e.g., Solidity). (**E**) Over 100 shape descriptors were evaluated in terms of how much they change when shapes are transformed (‘sensitivity’).

Although real-world objects are 3D, humans can make many inferences from 2D contours (e.g., [13, 26, 27]). Many 2D shape representations have been proposed—both for computational purposes and as models of human perception—each summarizing the shape boundary or its interior in different ways (**Figure 1B**; [24]). These include (but are not limited to) *basic shape descriptors* (e.g., area, perimeter, solidity; [28]), *local comparisons* (e.g., Euclidean distance; Intersection-over-Union, IoU; [29]), *correspondence-based metrics* (e.g., shape context; [30]), *curvature-based metrics* [31], *shape signatures* (see [24]), *shape skeletons* [32], and *Fourier descriptors* [33]. These different shape descriptors have complementary strengths and weaknesses.

Each one is sensitive to certain aspects of shape, but relatively insensitive to others (**Figure 1CDE**). We reasoned that by combining multiple shape descriptors into a multidimensional representation, we could create a composite shape metric that would capture the many different ways that human observers compare shapes.

### Complementary nature of different shape descriptors

To appreciate the complementary nature of different metrics—and the necessity of combining them—consider that human visual shape representation is subject to two competing constraints. On the one hand, to achieve stable object recognition across changes in viewpoint and object pose, it is useful for shape descriptors to deliver consistent descriptions across large changes in the retinal image (‘robustness’). On the other hand, to discriminate finely between different objects with similar shapes, shape descriptors must discern subtle changes in shape (‘sensitivity’). Different descriptors trade off these constraints, which is evident when organizing them along a continuum that describes their robustness to changes in shape across transformations—such as rotation, scaling, shearing, or adding noise.

We illustrate this for two transformations: rotation and bloating (**Figure 1CD**). Specifically, we transformed one exemplar from each of 20 different animal categories (e.g., birds, cows, horses, tortoise) with bloating and rotation transformations of varying magnitudes (see **Methods: Sensitivity/robustness analysis to transformation**). We find that the different descriptors are differentially sensitive to the transformations. Some shape descriptors (e.g., *solidity* which measures the proportion of the convex hull that is filled by the shape; [34]; **Figure 1CD**) are entirely invariant across rotations, while others (e.g., *shape context* which builds a histogram for each point on a shape and summarizes the angle and distance with all other points; [30]) are sensitive to object orientation. Yet descriptors invariant to rotation may be highly sensitive to other transformations, like bloating (**Figure 1CE**). Similarly, adding noise to a shape’s contour strongly affects curvature-based metrics, while only weakly affecting area or compactness (**Figure S1**). In **Figure 1E**, we plot how sensitive 109 different shape descriptors are to the changes introduced by rotation and bloating, highlighting the descriptors identified in **Figure 1B**. Interestingly, for these transformations, there is a tradeoff in sensitivity such that descriptors that are highly sensitive to bloating (e.g., *solidity*) tend to be less sensitive to rotation, and vice versa (e.g., *shape context*). In other words, different shape features have complementary strengths and weaknesses. More generally, the plot shows the wide range of sensitivities across different shape metrics, indicating that depending on the context or goal, different shape features may be more or less appropriate.

The key idea motivating our model is that human vision may resolve the conflicting demands of robustness and sensitivity by representing shape in a multidimensional space defined by many shape descriptors (**Figure 1B**). We do not intend the model to be a simulation of brain processes; i.e., it is unlikely to compute the specific model features considered here, (most of which are taken from previous literature; see Supplemental **Table S1**). Indeed, there are infinitely many other shape descriptors that could also be considered. Rather, we see the model as a proof of principle that human shape similarity can be predicted by representing shape using many, complementary geometrical properties.

To quantify how such features span the space of natural shapes, we evaluated >100 shape descriptors on >25,000 natural animal shapes and performed dimensionality reduction to derive a composite model. We then performed a series of experiments with naïve observers, to test the extent to which the model predicts human shape similarity judgments.

## Results

### Analysis of natural shapes

Different shape descriptors are measured in different units, so to combine the features into a consistent multidimensional space requires identifying a common scale. Given the importance of natural stimuli for human behavior, we reasoned that the relative scaling of the many feature dimensions likely reflects the distribution of feature values across natural shapes. We therefore assembled a database of over 25,000 animal silhouettes and for each of them measured a large set of shape descriptors (**Methods: Natural shape analysis**). For every pair of shapes, we computed the distances between each descriptor (scaled by their largest distance across the whole animal dataset; **Figure 1B**) and then combined the features into a single metric, yielding a multidimensional space. This space exhibited a prominent shape-based organization with nearby locations sharing similar shape characteristics. For example, approximately elliptical animals like rabbits, fish, and turtles lie near together (bottom left of **Figure 2A**), while spindly thin-legged shapes (e.g., spiders; see insets in **Figure 2A**) are found in the opposite corner of the space.

**Figure 2.**
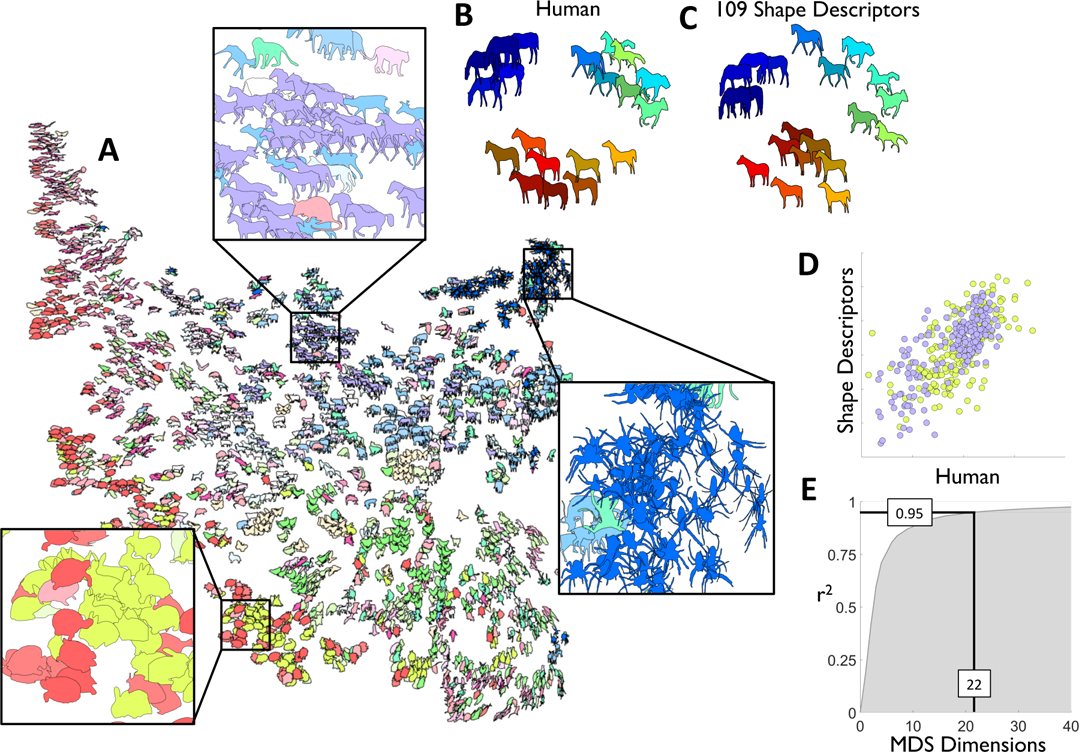
The high-dimensionality of natural shapes. (**A**) t-SNE visualization of 2000 animal silhouettes arranged by their similarities according to a combination of 109 shape descriptors. Colour indicates basic level category. Insets highlight local structure: bloated shapes with tiny limbs (left); legged rectangular shapes (middle); small spiky shapes (right). To test whether human shape similarity is predicted by in the high-dimensional animal space, we gathered human shape similarity judgments on horses (purple) and rabbits (yellow). (**B**) Human similarity arrangements (*n* = 15) of horse silhouettes (multidimensional scaling; dissimilarity: distances, criterion: metric stress). (**C**) Similarity arrangement for horse silhouettes in the full model based on 109 shape descriptors (multidimensional scaling; dissimilarity: distances, criterion: metric stress). Shapes with same colour across B and C are also the same. (**D**). Human arrangements correlate with the model for horse (purple) and rabbit (yellow) silhouettes (*r* = 0.69, *p* < 0.01). (**E**). Across 25,712 animal shapes, 22 dimensions account for >95% of the variance (multidimensional scaling; dissimilarity: distances, criterion: metric stress).

As an initial indicator of how well the features account for perceptual similarity with familiar objects, we took a subset of animal shapes, and measured human similarity judgements (**Figure 2B**) using a multi-arrangement method [35]. We find that the mean perceived similarity relationships between shapes were quite well predicted by distance in this feature space (**Figure 2CD**, *r* = 0.69, *p* < 0.01) suggesting that the 109 shape descriptors explain a substantial portion of the variance in human shape similarity of familiar objects. We suggest that at least some of the remaining variance is likely to be due to using familiar objects, for which high-level semantic interpretations are known to influence similarity judgments [36–40]—here, the perceived classes to which the animals belong, rather than their pure geometrical attributes.

We also find that many of the shape descriptors correlate with one another, yielding 22 clusters of related features (using affinity propagation clustering; [41]). Using Multidimensional Scaling across the 25,712 animal shape samples, we find that 22 dimensions account for more than 95.05% of the variance (**Figure 2E**), whereas the first dimension accounts for only 18.54% of the variance. Thus, while not all 109 shape descriptors are independent, a multidimensional space is indeed required to capture the variability inherent in animal shapes. We refer to this reduced 22-D space as ShapeComp (**Figure 1B**), and it is this model that forms the basis of the majority of our subsequent analyses.

ShapeComp’s dimensions are composites (i.e., weighted linear combination) of the original shape descriptors, which makes the model fully interpretable, unlike other model classes, (e.g., neural networks, whose inner functioning researchers still struggle to interpret [42, 43]). Although we do not believe the brain explicitly computes these specific dimensions, they do organize novel shapes systematically (**Figure S2**), and also tend to re-appear in different random subsets of animal shapes or combination of shape descriptors (**Figure S3**). However, because MDS creates a rotation invariant space, individual dimensions should not be thought of as perceptually meaningful ‘cardinal axes’ of shape space. Rather it is the space as a whole that describes systematic relationships between shapes.

### Creating a ‘shape space’ for novel naturalistic outlines

To reduce the impact of semantics on shape similarity judgments, we next created novel (unfamiliar) shapes using a Generative Adversarial Network (GAN) trained on the animal silhouette database (see **Methods**). GANs are unsupervised machine learning systems that pit two neural networks against each other (**Figure S4A**), yielding complex, naturalistic, yet largely unfamiliar novel shapes. The GAN also allows parametric shape variations and interpolations in a continuous ‘shape space’ (**Figure S4BCD**). Overall, the GAN shapes appear ‘object-like’ (**Figure S5A**), but observers agree less about their semantic interpretation, compared with animal shapes (**Figure S5**), making them better stimuli for assessing pure shape similarity. We next sought to test more rigorously (a) whether distance in ShapeComp space predicts human shape similarity, (b) whether ShapeComp provides information above and beyond simpler metrics like pixel similarity, (c) whether human shape similarity is indeed multidimensional, and (d) whether ShapeComp identifies perceptual nonlinearities in shape sets.

### Distances in ShapeComp model predict human shape similarities

A key criterion for any perceptual shape metric is that pairs of shapes that are close in the space (**Figure 3A, top**) should appear more similar than pairs that are distant from each other (**Figure 3A**, **bottom**). To test this, we generated 250 pairs of novel GAN shapes, ranging in their ShapeComp distance (i.e., predicted similarity), and asked 14 participants to rate how perceptually similar each shape pair appeared (**Figure 3B**). We find that distance in ShapeComp correlates almost perfectly with the mean ratings across observers (*r*^2^ = 0.99, *p*<0.01) showing that ShapeComp predicts human shape similarity very well for novel unfamiliar 2D shapes.

**Figure 3.**
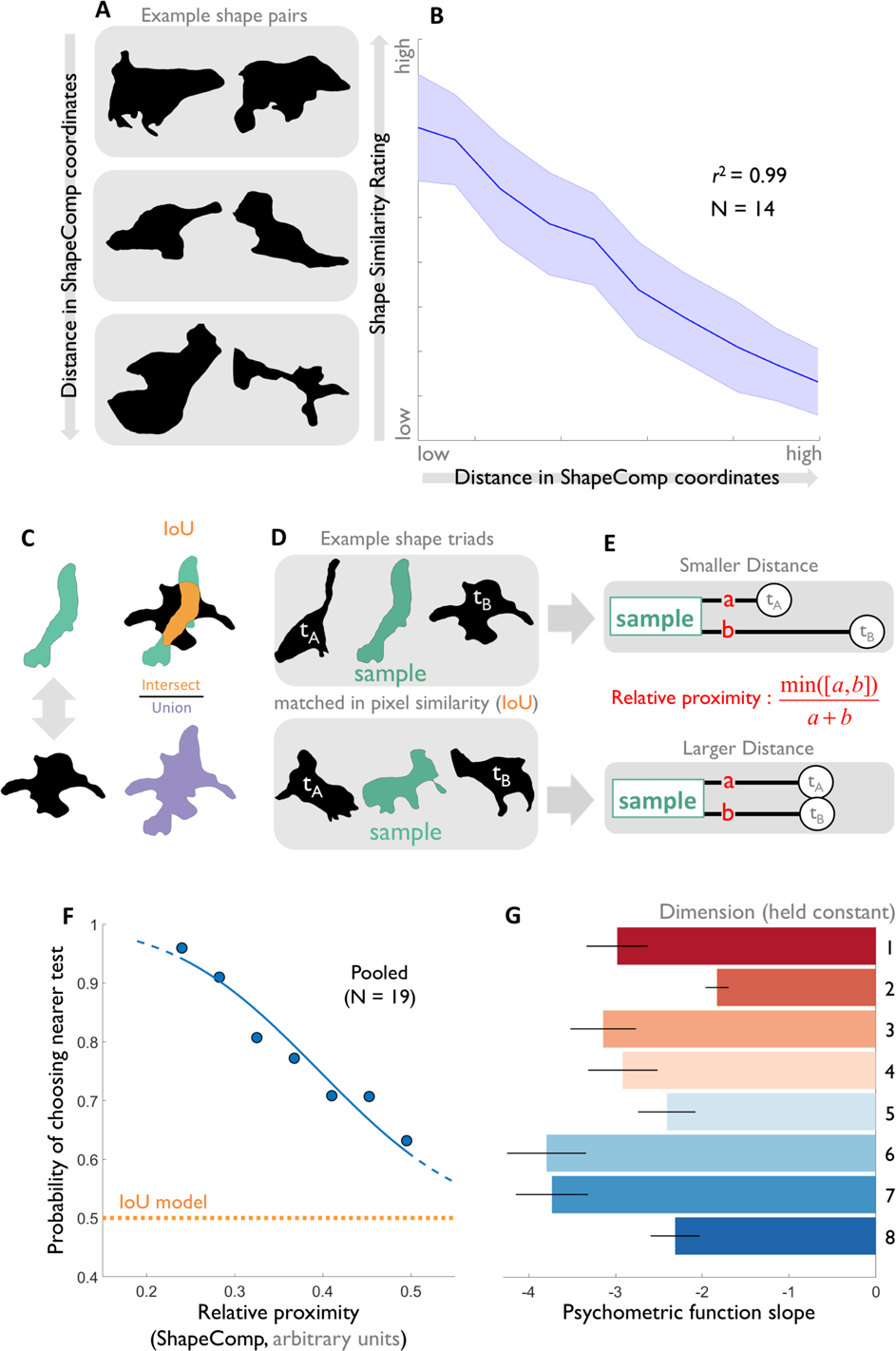
ShapeComp predicts human shape similarity across pairs of shapes. (A) Example shape pairs that varied as a function of ShapeComp distance. **(B)** Shape similarity ratings from 14 observers correlate almost perfectly with distance in ShapeComp’s 22-dimensional space. **(C)** Pixel similarity was defined as the standard Intersection-over-Union (IoU; [29])**(D)** Observers viewed shape triads and judged which test appeared more similar to the sample. (**E**) ShapeComp distance between test and sample were parametrically varied but pixel similarity was held constant. (**F**) Mean probability across participants, that the closer of two test stimuli was perceived as more similar to the sample, as a function of the relative proximity of the closer test shape. Blue: psychometric function fit; orange: prediction of IoU model. (**G**) Results of experiment in which distances from test to sample were equated for one ShapeComp dimension at a time. Mean psychometric functions slopes were much steeper than predicted if observers relied only on the respective dimension.

**Figure 4.**
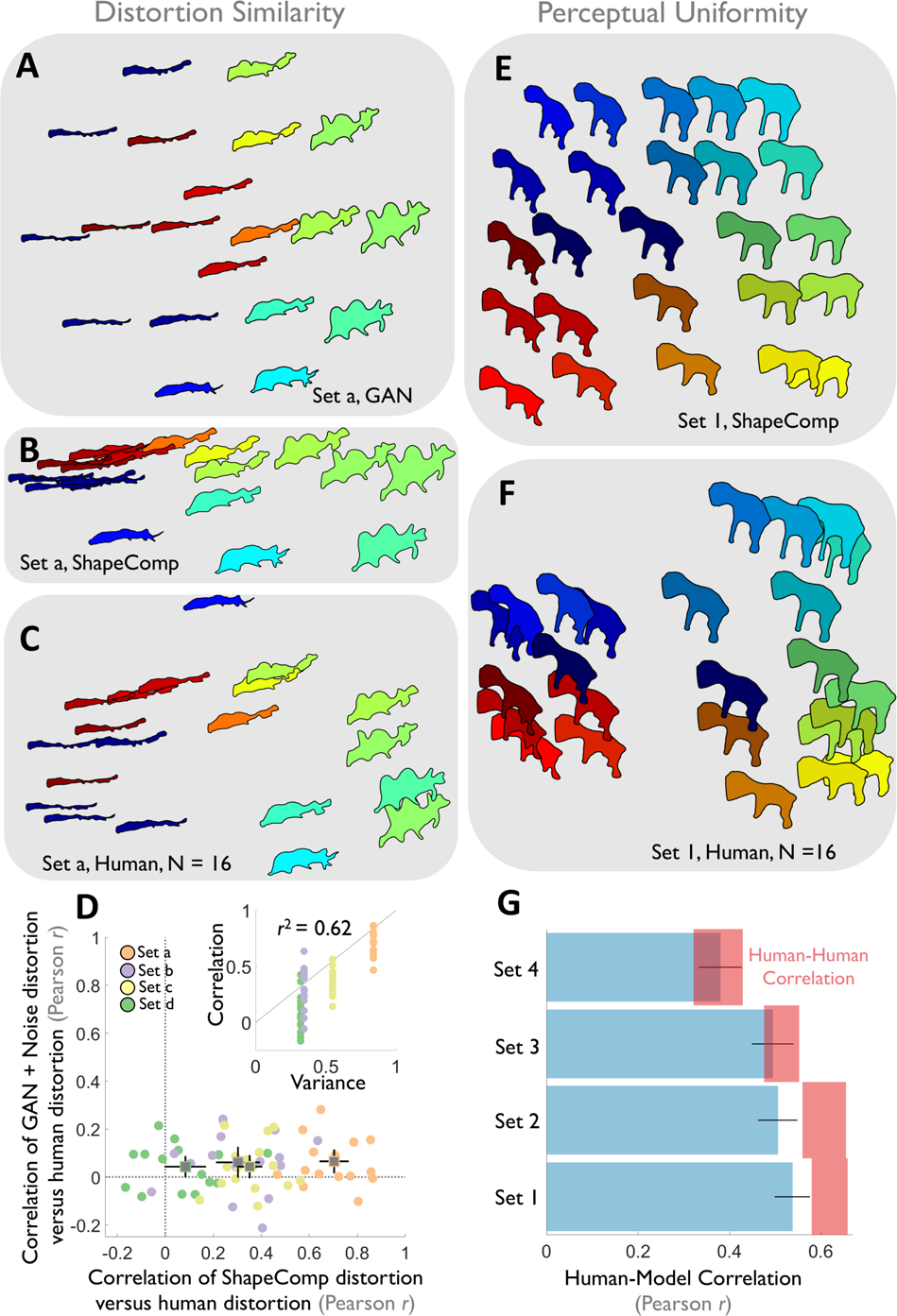
ShapeComp predicts human shape similarity in shape arrays. **(A)** Example 2D shape array that is uniform in GAN space but **(B)** distorted by ShapeComp is similarly **(C)** distorted by humans (left; mean across 16 participants). Across arrangements, shapes with same colour are also the same. **(D)** Non-uniformities for individual participants (dots) in 4 shape sets (colours) selected to vary slightly in their GAN and ShapeComp predictions (Pearson correlation coefficient: Set a, *r* = 0.59; Set b, *r* = 0.74; Set c, *r* = 0.75; Set d, *r* = 0.68). Squares show average across subjects for given set, where error bars show ± 2 standard errors. ShapeComp accounted for perceptual distortions away from the original GAN coordinates better than GAN+noise model. Inset: Correlation of ShapeComp distortion with human distortion as a function of cumulated variance in shape set across ShapeComp dimensions. Human distortions better line up with ShapeComp when there is more variance across shape sets as predicted by ShapeComp. Grey reference line shows y=x. **(E)** Example 2D shape array that is roughly uniform in ShapeComp and highly correlated to the GAN arrangement (r>0.9). **(F)** Mean arrangement by 16 human observers. (**G**) In 4 shape sets that are highly correlated in terms of GAN and ShapeComp arrangements, human responses are nearly indistinguishable from the predictions of ShapeComp (blue), given the inherent noise across observers measured as the lower noise ceiling (red; correlation of each participant’s data with mean of others).

### Distance in ShapeComp goes above and beyond pixel similarity

A standard way to measure the physical similarity between shapes is the Intersection-over-Union quotient (IoU; [29]; **Figure 4C**). For similar shapes, the area of intersection is a significant proportion of the union, yielding IoU values approaching 1. In contrast, when shapes differ substantially, the union is much larger than the overlap, so IoU approaches 0.

To test whether human shape similarity can be approximated by such a simple pixel similarity metric or rather relies on more sophisticated mid-level features like those in ShapeComp, we created stimulus triplets, consisting of a *sample* shape, plus two *test* shapes, which were equally different from the sample shape in terms of IoU but which differed in ShapeComp distances (**Figure 3D** and **Methods: pixel similarity triplets**).

This allowed us to isolate the extent to which ShapeComp predicted additional components of shape similarity, above and beyond pixel similarity. The magnitude of the difference between tests and sample in ShapeComp was varied parametrically across triplets, so that sometimes one test was much nearer to the sample than another test (**Figure 3E**). Nineteen new participants viewed the triplets and were asked which of the two test shapes most resembled the sample on each trial. If shape perception is perfectly captured by IoU, the two test stimuli should appear equally similar to the standard, yielding random responses (**Figure 3F** orange line). However, we find that the slope of a psychometric function fitted to the observers’ judgments is significantly steeper than zero (**Figure 3F** blue line; *t* = −32.3, df = 18, *p* <0.01). This indicates that ShapeComp correctly predicts which of the two shapes was more similar to the standard even when pixel similarity is held constant. This shows that human shape similarity relies on more sophisticated features than pixel similarity alone.

### Multidimensionality of human shape similarity

We suggested that human shape similarity judgments are inherently multidimensional, as observers consider multiple aspects of shape when comparing two stimuli. To test this, we generated triplets in which the test shapes were equated to a given sample shape in terms of one of ShapeComp’s 22 dimensions but varied in terms of the remaining dimensions. The same nineteen participants as in the pixel similarity experiment were shown these triplets and again reported which test shape appeared most similar to the sample. If shape perception is entirely captured by any single dimension, the two test stimuli should appear equally similar to the sample, yielding random responses. Yet **Figure 3G** shows that fitted psychometric function slopes were significantly steeper than zero. This indicates that human shape perception is inherently multidimensional—when each dimension was held constant, the variations in the remaining dimensions dominated perception.

We also re-analyzed the ratings from **Figure 3B**, comparing the human judgments to each ShapeComp dimension. Each dimension on its own accounted for only a small portion of the variance (**Figure S6**), again indicating the inherently multidimensional nature of human shape similarity judgments. Interestingly, the fall-off in variance in natural shapes (in **Figure 2E**) closely tracks the variance explained by each ShapeComp dimensions (*r^2^* = 0.81, p<0.01). This suggests that the dimensions underlying human shape perception are weighted by the extent to which they capture variations between natural shapes. Together, these results show that human shape similarity relies on multiple ShapeComp dimensions—highlighting the importance of combining many complementary shape descriptors into ShapeComp—and suggest that shape similarity is tuned to the structure of natural shapes.

### Identifying perceptual nonlinearities in shape spaces of novel objects

An important test for any human shape similarity metric is its ability to predict similarity relationships within arrays of multiple shapes. To assess this, we tested how well ShapeComp identified perceptual non-uniformities in shape spaces generated with the animal-trained GAN. **Figure 4A** shows an example 2D GAN shape array sampled at uniform radial distances that ShapeComp’s prediction (in 2D) is perceptually non-uniform (**Figure 4B**). Using a multi-arrangement task (**Methods**), we find that human perceived similarities within these arrays resembled the nonuniformities predicted by ShapeComp (mean responses from 16 participants: **Figure 4C**).

Are ShapeComp’s perceptual distortions from the original uniform GAN space better at predicting shape similarity than a random model? To measure distortions between shape arrays, we computed differences between two similarity matrices — each standardized to have unit variance — where larger differences lead to larger distortions. To test whether ShapeComp is better than a random model, we developed a GAN+noise model that distorts the original GAN space by adding random Gaussian perturbations to the original GAN latent vector coordinates. We set the noise level of the model to maximize its chance of accounting for the human distortions by matching the overall distance of the noise perturbations from the original GAN space with the overall perturbations of the human observers (from the original GAN space). Across 4 different shape sets where GAN and ShapeComp spaces tended to be less correlated with one another (0.59<r<0.75), perceptual distortions in GAN space by individual observers were better accounted for by ShapeComp than the GAN+noise model (**Figure 4D**). Further, the overall variance in ShapeComp’s coordinates across a shape set was highly predictive of how well ShapeComp distortions matched humans: Greater variance in ShapeComp lead to more overlap with humans (*r^2^* = 0.7, *p*<0.01; **Figure 4D** inset). Thus, ShapeComp correctly predicted the direction of perceptual nonlinearities in the GAN space. This is striking given that the GAN arrays and ShapeComp are highly correlated, and thus already share much of the variation across their arrangements of the shape sets.

### Deriving perceptually uniform shape spaces of novel objects

With the ability to measure perceptual non-uniformities in hand, ShapeComp can also be used to create perceptually uniform arrays of novel objects. To do this, we searched for uniform arrays the GAN’s latent vector representation that were highly correlated with ShapeComp (r>0.9). **Figure 4E** shows an array that ShapeComp predicts should be arranged almost uniformly. Human similarity arrangements (mean response from 16 participants; **Figure 4F**) are consistent with ShapeComp in terms of the relative ordering of the shapes. Across 4 different shapes sets, human responses are nearly indistinguishable from the predictions of ShapeComp, given the inherent noise across observers (**Figure 4G**). Thus, by combining the high-dimensional outputs of the GAN with ShapeComp, we can now automatically create a large number of perceptually uniform shape spaces.

## Discussion

Many previous studies have sought to measure shape similarity for both familiar and unfamiliar objects [38, 40, 44–53]. Despite this, the representation of shape in the human visual system remains elusive, and the basis for shape similarity judgments remains unclear. In part, this is due to the numerous potential shape descriptors proposed in the past, including simple metrics, like solidity [28], and contour curvature [31], and more complex metrics like shape context [30], part-based ones [1], Fourier descriptors [33,45,54], radial frequency components [46], and shape skeletons [32,51,52,56], and models based on generalized cylinders for describing 3D animal-like objects [57].

Which of these features does the brain use to represent and compare shapes? We suggest the brain combines many different features into a multidimensional space, allowing the visual system to resolve the conflicting constraints of *sensitivity* and *robustness* to transformations. Indeed, we suggest that the precise feature set is less important than the space spanned by the features. Given the multiplicity of cells that contribute to representations of shapes and objects in ventral processing stream, it may not even be possible to describe a complete and unique set of features that the human visual system uses. In fact, the response properties of cell populations may vary significantly across observers, yet similarity relationships between shapes could be preserved. Hence it makes more sense to focus on the feature space as a whole, rather than the contributions of individual putative dimensions.

Multidimensional representations allow subsequent visual processes to selectively attend to different aspects of shape, optimizing features for environmental statistics and task demands [58–61]. For example, Morgenstern, Schmidt, and Fleming [51] showed that in one-shot categorization observers tend to base their judgments of whether two novel objects belong to the same category on different features depending on the specific shapes to be compared. Because the features weights in ShapeComp are derived from the statistics of animal shapes, it is well suited to distinguishing natural shapes. It is intriguing that no fitting was necessary to predict human shape similarity judgments using ShapeComp — the raw weights derived from ca. 25,000 natural silhouettes account for most of the variance in **Figure 3B**. This suggests that natural shape statistics may play a central role in determining the space humans use to represent and compare shapes. Indeed, ShapeComp also correctly predicts previous shape similarity data. For example, Li et al. [53] constructed a ‘perceptually circular’ stimulus set, which ShapeComp predicts quite well (**Figure S7**).

Paired with a GAN trained on animal silhouettes ShapeComp also provides a useful tool for automating the analysis and synthesis of complex naturalistic 2D shapes for future experiments in cognitive psychology and neuroscience. Novel, perceptually-uniform stimulus arrays can be generated and probed on the fly (**Figure 4**, S8), for example, adaptively modifying stimuli in response to brain activity during an experiment.

ShapeComp can also help create single- or multi-dimensional arrays (**Figure S8A-C**), or stimulus sets that are perceptually equidistant from a given probe stimulus (**Figure S8D**). Once stimulus sets are controlled for image-based properties, the role of higher-level aspects of object representations can be probed in perception, visual search, memory, and other tasks.

### Limitations

There are a number of respects in which ShapeComp could be improved in further work. First, for many applications it would be desirable to characterize similarity in 3D (e.g., computer vision and computer graphics [25]; video analysis [62]; topology mapping [63]; molecular biology [21]). Second, shapes in the natural world are often occluded, while ShapeComp was trained only on non-occluded shapes. Third, ShapeComp was trained only on animal shapes. While the training set spans a very wide range of shape characteristics, future studies could refine ShapeComp by covering other major superordinate categories such as plants, furniture, tools and vehicles. This would probably modify the weighting of individual dimensions of ShapeComp, yet may further improve ShapeComp’s predictions of human similarity judgments. Fourth, while ShapeComp pools 109 different descriptors from across the literature, there are many others that were not included. Incorporating additional features would likely change the precise estimates of similarity made by ShapeComp (although, **Figure S3** suggests that using different subsets of features yields similar composite dimensions in MDS). Yet, we believe that there is no one single shape descriptor that perfectly captures all of human shape similarity perception, and that the general approach of pooling multiple descriptors provides robust and sensitive representations. Fifth, as a model of human perception, ShapeComp is entirely parameter-free in the sense that no fitting was used to adjust the features or their weights to improve predictions of human judgments. We saw this as an important component of testing whether weightings derived from natural shapes predict human perception. However, with over 100 features, ShapeComp’s predictions could almost certainly be further improved by explicitly fitting to human data. Finally, ShapeComp is not a physiologically plausible model of shape representation processes in the human brain. Future research should seek to model in detail the specific features in the neural processing hierarchy that represent shapes in a multidimensional space [64]. We believe that paired with novel image-generating methods, like GAN’s ShapeComp can play a central role in mapping out visual shape representations in cortex.

Shape can be described in many different ways, which have complementary strengths and weaknesses. We have shown that the brain takes advantage of this by combining many different shape descriptors into a multidimensional representation that is tuned to the statistics of natural objects. The ShapeComp model correctly predicts human shape perception across a wide range of conditions. It captures perceptual subtleties that conventional pixel-based metrics cannot, and provides a powerful tool for generating and analysing stimuli. Thus, ShapeComp not only provides a benchmark for future work on object perception, but also explains how human shape processing is simultaneously sensitive, robust and flexible.

## Materials and Methods

### Natural shape analysis

For 25,712 animal shapes—purchased from shutterstock, or gathered from previous work (e.g., [65])—we calculated 109 shape descriptors thought to be important for recognition, synthesis, and perception [24]. The shapes’ x,y coordinates (382×2 resolution) were uniformly scaled to {0-1}. For shape descriptors requiring an initial shape point, this was set to the point with the smallest x-value.

### Shape Descriptors

Shape Descriptors consisted of simple descriptors like area and perimeter, to more complex descriptors like the shape skeleton. A full list of the 109 descriptors is available in Supplemental **Table S1**.

### Multidimensional scaling and ShapeComp model

We used classical MDS to find an orthogonal set of shape dimensions that captures the variance in the animal dataset. For each shape descriptor, we computed the Euclidean distance, for each pair of shapes in the dataset:

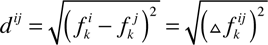

where *d^ij^* is the distance between stimulus *i* and *j* on shape descriptor *k* and *f ^i^_k_* and *f ^j^_k_* are the values on shape descriptor *k* for stimuli *i* and j. Once the computation for all pairwise comparisons was complete, the distances were assembled into a 25,712×25,712 similarity matrix and normalized by their largest distance. We computed this normalized distance,

*d*^^^, for all shapes and shape descriptors to form 25,712×25,712×109 entry matrix (shapes^2^ × shape descriptors). We then computed a 109-dimensional Euclidean distance *D* across the shape descriptors for shape pair *i* and *j*, as follows

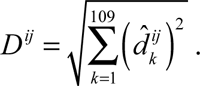

We then computed classical MDS on the resultant 25,712×25,712 similarity matrix.

### Estimating coordinates for new shapes in pre-existing shape spaces

We estimate the coordinates for a new shape in the high-dimensional animal MDS space by comparing the shape descriptors for the new shape with a subset of animal shapes, computing a new MDS solution, and then moving this new MDS solution using Procrustes to the high-dimensional animal MDS space. Specifically, we computed the Euclidean distance between the new shape and 500 shapes already located in the animal space, to assemble a 501×501 similarity matrix, and scaled by the largest distance for each feature distance in the complete animal dataset. We did this for all shape descriptors to form a 501×501×109 matrix (shapes^2^ × shape descriptors). We then computed the 109-dimensional Euclidean distance *D* across shape descriptors yielding a 501×501 similarity matrix. Applying Classical MDS produced a new coordinate space for the original 500 shapes. We used procrustes to transform the MDS coordinates for the 500 animal shapes from the new coordinate space to the original coordinate space. We then applied this transformation to the new shape to move it into the original shape space.

### GAN shapes

A Generative Adversarial Network, GAN [66, 67], was trained using MatConvNet in MATLAB to synthesize shapes that it could not distinguish from the animals shapes database. The network architecture and hyperparameters were the same as in Radford et al. [67], except for the following. The latent *z* vector was 25×1 (rather than 100×1) and one of the dimensions of the remaining filter sizes was reduced (from initially matching the other dimension) to 2. A series of four “fractionally-strided” convolutions then converted the latent vector’s high-level representation into the shapes’ spatial coordinates. After 106 training epochs, we generated novel GAN shapes by inputting random vectors into the GAN latent variable. We blurred the shapes with a Gaussian filter limited to two neighbouring contour points and selected shapes without self-intersections.

### ShapeComp22 Network

We trained a convolutional neural network using MATLAB’s neural network toolbox to take as input a 384×2 contour through 3 convolution neural layers and output the 22-dimensional MDS coordinate. To do this, we created a training set of 1,000,000 GAN shapes (800,000 training, 200,000 test images) and then computed their 22D MDS coordinates (see *Estimating coordinates for new shapes in pre-existing shape spaces)*. These coordinates served as the desired network output. The network architecture and training hyperparameters are shown in **Figure S9**. Novel input shapes yield an estimate of the 22D MDS coordinate as output. (See also **Supplemental Discussion** on **ShapeComp22 Network**)

### Sensitivity/robustness analysis to transformation

We transformed one sample from each of the 20 animal categories (from [65]) with 2D transformations (e.g., rotation, shear, bloating, noise) of varying strengths. For each sample, we examined how sensitive each shape descriptor was to a given transformations by computing differences between shape descriptors for the original shape with the transformed version. We then scaled these differences to range from 0 – 1, and took the mean. We then took the nanmean across the 20 samples as the sensitivity to a given transformation, where larger values indicate less sensitivity.

### General experimental methods

#### Participants

Participants, paid at a rate of 8 euros per hour, signed an informed consent approved by the ethics board at Justus-Liebig-University Giessen and in accordance with the Code of Ethics of the World Medical Association (Declaration of Helsinki). Participants reported normal or corrected-to-normal vision.

#### Procedure

All experiments were run with an Eizo ColorEdge CG277 LCD monitor (68 cm display size; 1920 x 1200 resolution) on a Mac Mini 2012 2.3 GHz Intel Core i7 with the psychophysics toolbox [68, 69] in MATLAB version 2015a. Observers sat 57cm from the monitor such that 1 cm on screen subtended 1° visual angle.

#### Shape Similarity in Stimulus Pairs/Triplets

##### Participants

14 observers participated in the shape similarity rating experiments and 19 different observers participated in the *pixel similarity* and *ShapeComp22 dimensions* experiment. Mean age was 24.3 (range: 20-33).

##### Pairwise similarity ratings

250 GAN shape pairs were chosen that spanned a large range of distances in ShapeComp. On each trial, stimuli were shown side by side and observer adjusted a slider to indicate similarity ratings from 0-100 using the mouse. Shapes subtended ∼15°. Shape position (right or left side) was randomized on each trial. Shape pairs were presented in random order.

##### Pixel similarity triplets

Using the Genetic Algorithm in MATLAB’s Global Optimization toolbox, we used the ShapeComp22 network to find shape triplets in which a *sample* shape varied in its ShapeComp distance from two *test* shapes, *t_A_* and *t_B_*, while maintaining the same pixel similarity to both. Specifically, we computed the ShapeComp distance from the *sample* to each *test*, *a* for *t_A_* and *b* for *t_B_* (**Figure 3E**). We then represented the distances from these test shapes to the sample as a ratio between the smaller of the distances to the sum of their distances:

*min(a, b*) / *(a + b*)

Small values of this ratio indicate one *test* was much closer to the *sample* shape in terms of ShapeComp. A maximum value of 0.5 indicates both tests are equally far from the sample. 70 triplets were created and binned into 7 bins ranging 0.2–0.5, where each bin contained ∼10 triplets. On each trial, the *sample* shape was presented centrally, flanked by two *test* shapes (whose position, left or right of *sample* was randomized). Shapes subtended 12°. Pixel similarity, held constant between the *sample* and the *test* shapes, was defined as the Jaccard index (1 - intersection-over-union; [29]). High values indicate high pixel similarity.

##### ShapeComp dimensions experiment

Similar to the pixel similarity experiment, using the Genetic Algorithm in MATLAB’s Global Optimization toolbox, we used the ShapeComp22 network to find shape triplets in which a *sample* shape varied in its ShapeComp distance from two *test* shapes, *t_A_* and *t_B_*, while maintaining the same value on one of the ShapeComp dimensions {1–8}. The distance between *sample* and *test* shapes was represented with the ratio described in the *pixel similarity triplets*.

#### Shape Similarity in shape arrays

##### Animal shape similarity

*Participants.* 15 observers participated (mean age: 24.7 years; range 20–35).

*Stimuli.* Two sets of twenty shapes (either rabbits or horses) from [66].

##### Perceptual non-uniformities

*Participants.* Two groups of 16 observers (mean age: 24.45 years; range: 18–41), including the first author who was the only author and participant in both groups. *Stimuli.* Four GAN shape sets were selected that ranged in their correlation with ShapeComp (0.59 < *r* < 0.75). One group of participants arranged two sets with 20 shapes (set a, *r* = 0.59; set b, *r* = 0.75). Another group arranged two sets with 25 shapes (set c, *r* = 0.68; set d; *r* = 0.74).

### Perceptually uniform shape spaces

*Participants.* Two groups of 16 observers (mean age: 25.03 years; range: 18–41), including the first author who was the only author and participant in both groups. *Stimuli.* Four sets of 25 shapes for which GAN and ShapeComp predicted similar pairwise distances (*r* > 0.9). One group of participants arranged two sets that were uniform in ShapeComp (set 1 and 2). Another group arranged two sets that were uniform in GAN space (set 3 and 4).

*Procedure.* Experiments were run in MATLAB using the multi-arrangement code provided by Kriegeskorte & Mur[35] and adapted for the Psychophysics Toolbox [68, 69]. On each trial, participants used the mouse to arrange all stimuli by their similarity relationships to one another within a circular arena. At the start of each trial, stimuli were arranged at regular angular intervals in random order around the arena. To the right of the arena, the current and last selected objects were shown larger in size (15°). Once an arrangement was complete, participants pressed the Return key to proceed to the next trial. The next trials showed a subset of the objects from the first trial based on the ‘*lift-the-weakest*’ algorithm [35]. The arrangements ended after 12 minutes had elapsed.

## Acknowledgments

This research was funded by the DFG funded Collaborative Research Center “Cardinal Mechanisms of Perception” (SFB-TRR 135) and the ERC Consolidator award ‘‘SHAPE’’ (ERC-CoG-2015-682859). We thank Saskia Honnefeller, Jasmin Kleis, and Marcel Schepko for their help setting up the experiments and running initial pilot studies.

## Competing Interests

The authors declare that there are no financial and non-financial competing interests.

## Supplemental Methods

### Category judgement experiment

*Participants.* Twenty participants classified GAN shapes, and another 20 classified animal shapes.

*Stimuli.* Photographs (9×12.5 cm) of 100 GAN shapes with no-self intersections (randomly selected from the GAN latent space) and 20 animal shapes from Bai et. al [1]. Each photograph had a number to indicate shape (1-100 for GAN shapes, 1-20 animal shapes).

*Procedure.* Experimenter shuffled the cards, and placed them in front of participant. Participant picked up the top card and placed it roughly arms length from their view. They called out the number on the card, and were then asked to judge the category of the shape on the card. Participants had the option of saying that the shape is does not appear like any known category. Experimenter entered the responses, while the participant picked up the next card from the pile. This process continued until the participant finished classifying the whole stack.

### Supplemental Discussion

#### A neural network that approximates ShapeComp

Although the features underlying ShapeComp are both image computable and interpretable, in practice, the codebase is convoluted—as it draws on many different sources—and the computation of all 109 features along with pairwise comparisons of the with 109 pre-computed features from a large dataset of stored animal shapes is too slow for real-time applications. Moreover, as argued above, individual features are less important than the space spanned by them in concert. Thus, to consolidate ShapeComp into a single, high-speed model, we trained a 3-layer convolutional neural network on 1,000,000 GAN shapes that spanned the high-dimensional space (**Methods:** *ShapeComp22 Network*). The network takes as input closed contours (with a resolution of 384×2) and outputs a 22-dimensional vector, representing the values of each of the dimensions of ShapeComp. **Figure S9** shows that the network is also highly related to human similarity judgements. The average error of the network in estimating ShapeComp is within the ShapeComp range that human observers tend to judge as very similar (an overall distance in ShapeComp <0.8), indicating that the neural network provides sufficiently good approximation to ShapeComp for most practical purposes.

More important than absolute deviation between ShapeComp coordinates is how ShapeComp captures the relationship between shapes; the network’s predicted distances across the upper triangular matrix of all pairwise combinations in 1000 untrained shapes is highly related to ShapeComp (*r*^2^ = 0.87, *p* <0.01). This allows experimenters to identify where arbitrary shapes lie within the 22D ShapeComp space making the network an easy way to measure similarity across arrays or pairs of shape, and paired with a shape generation tool (like GANs), allows the creation of perceptually uniform shape spaces.

### Supplemental Figures

**Figure S1.**
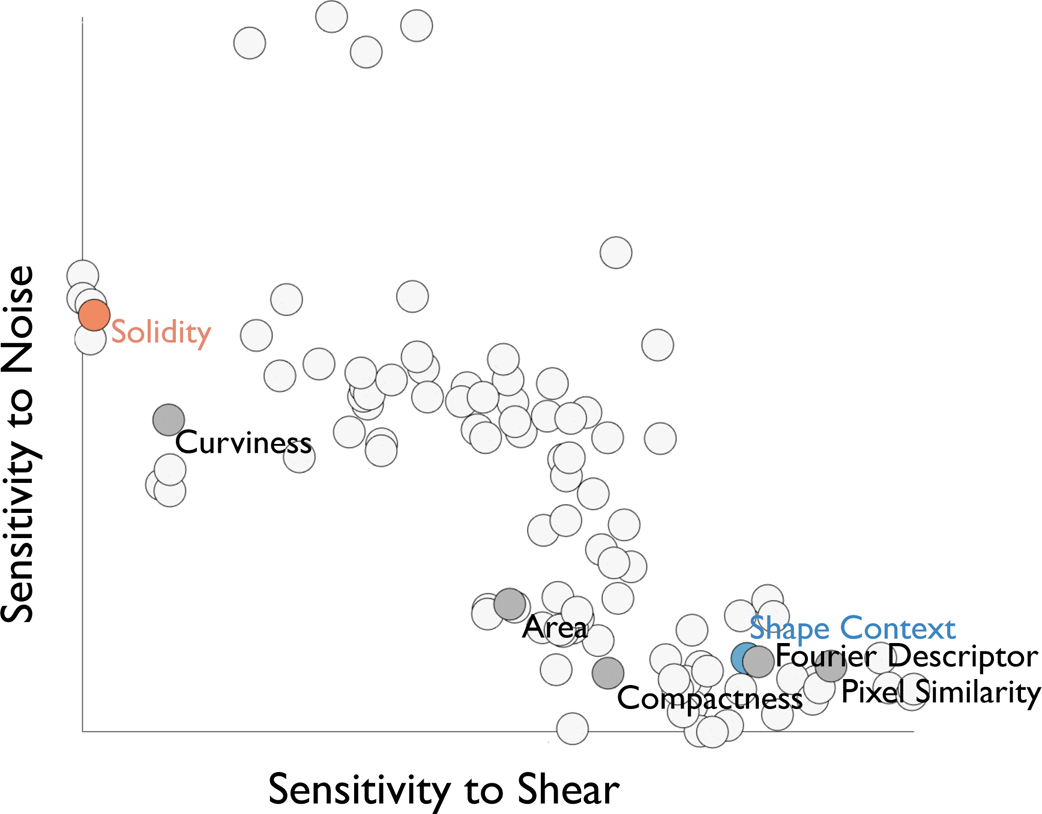
Over 100 shape descriptors evaluated in terms of their ‘sensitivity’, i.e., how much they changed when shapes were transformed by noise and shear. Here, the curvature signatures are more sensitive to noise than shear, while compactness is less sensitive to noise than shear. That different descriptors are tuned to different transformations highlights their complementary nature.

**Figure S2.**
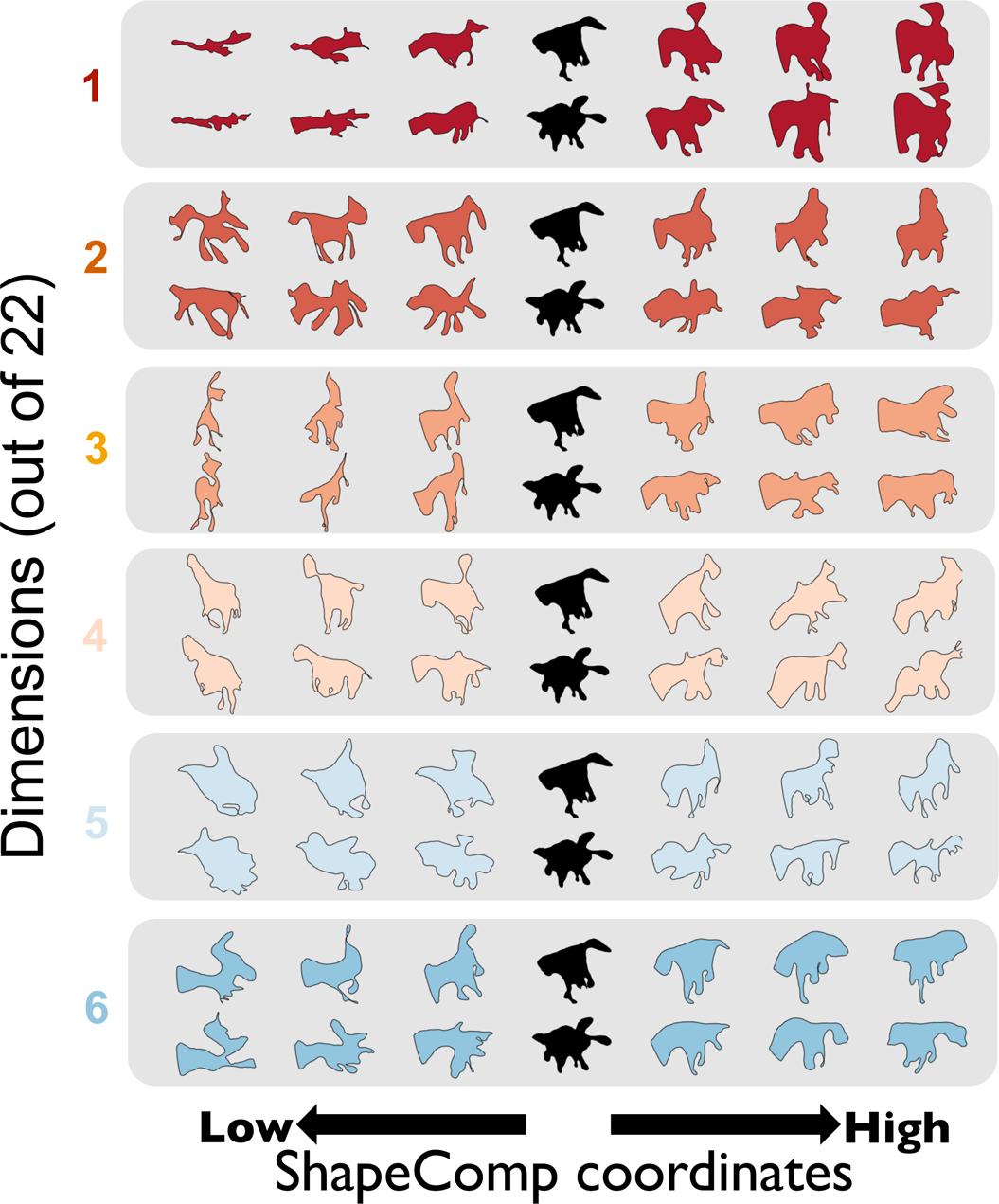
Example GAN shapes that vary along the first 6 MDS dimensions. Two shapes (in black) are varied along one dimension (in different colours, dimensions 1-6) while the remaining dimensions are held constant. The individual dimensions recovered by MDS should not be considered cardinal directions of the shape space as MDS is rotation invariant. Nevertheless, at least the first few dimensions are systematically organized with distinctive and different types of shape at opposite ends of each scale. However, much like the properties of receptive fields in mid- and high-level visual areas, it is not always easy to verbalize the properties underlying each MDS dimension: Dimensions 1 and 3 appear to modulate horizontal and vertical aspect ratio, respectively, but other factors like number and extent of limbs also vary. The different GAN shapes that varied in their MDS coordinates were optimized with a genetic algorithm from MATLAB’s global optimization toolbox to reduce RMS error between a GAN shapes 22-D representation and a desired 22-D representation.

**Figure S3.**
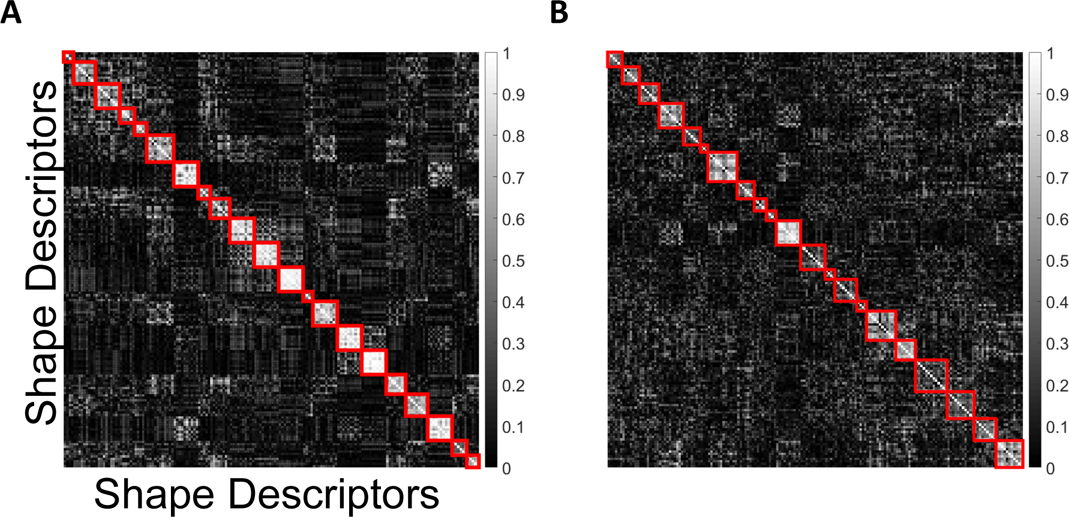
A high-dimensional features space for natural shapes. (A) We computed MDS across 10 different sets of 500 randomly chosen animal sets. We find correlated MDS dimensions across different random sets of 500 animal shapes, so that one dimension in set 1 is correlated to another dimension in the other 9 sets. **(B)** Across 10 different sets of shape descriptors (randomly selected 55 out of 109), we also find correlations across MDS dimensions.

**Figure S4.**
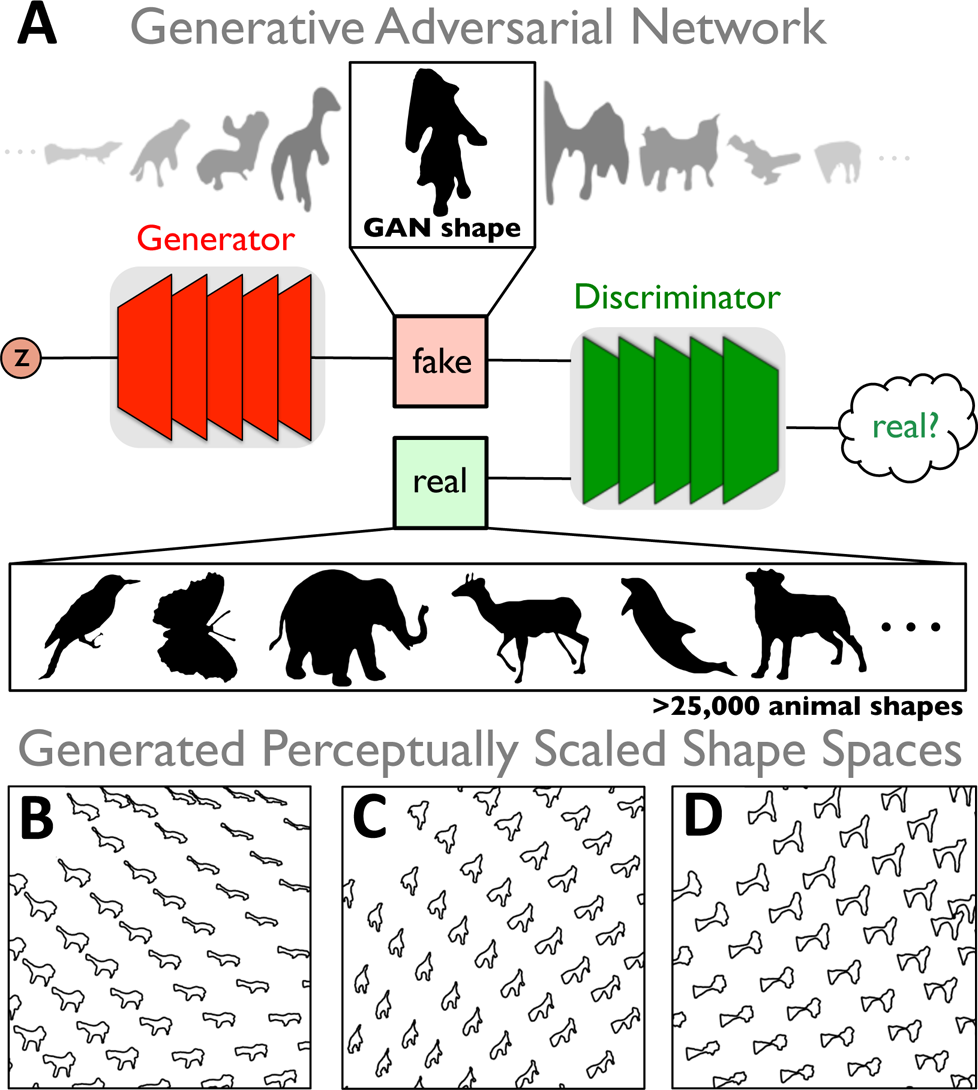
GANs produce novel naturalistic shapes. (A) Cartoon depiction of a Generative Adversarial Networks (GANs) that synthesizes novel shape silhouettes. GANs are unsupervised machine learning systems with two competing neural networks. The generator network synthesizes shapes, while the discrimantor network, distinguishes shapes produced by generator from a database of over 25,000 animal silhouettes. With training, the generator learns to map a high-dimensional latent vector ‘*z*’ to the natural animal shapes, producing novel shapes that the discrimantor thinks are real rather than synthesized. Systematically moving along the GANs high-dimensional latent vector produces novel shape variation and interpolations across a shape space **(B,C,** and **D)**.

**Figure S5.**
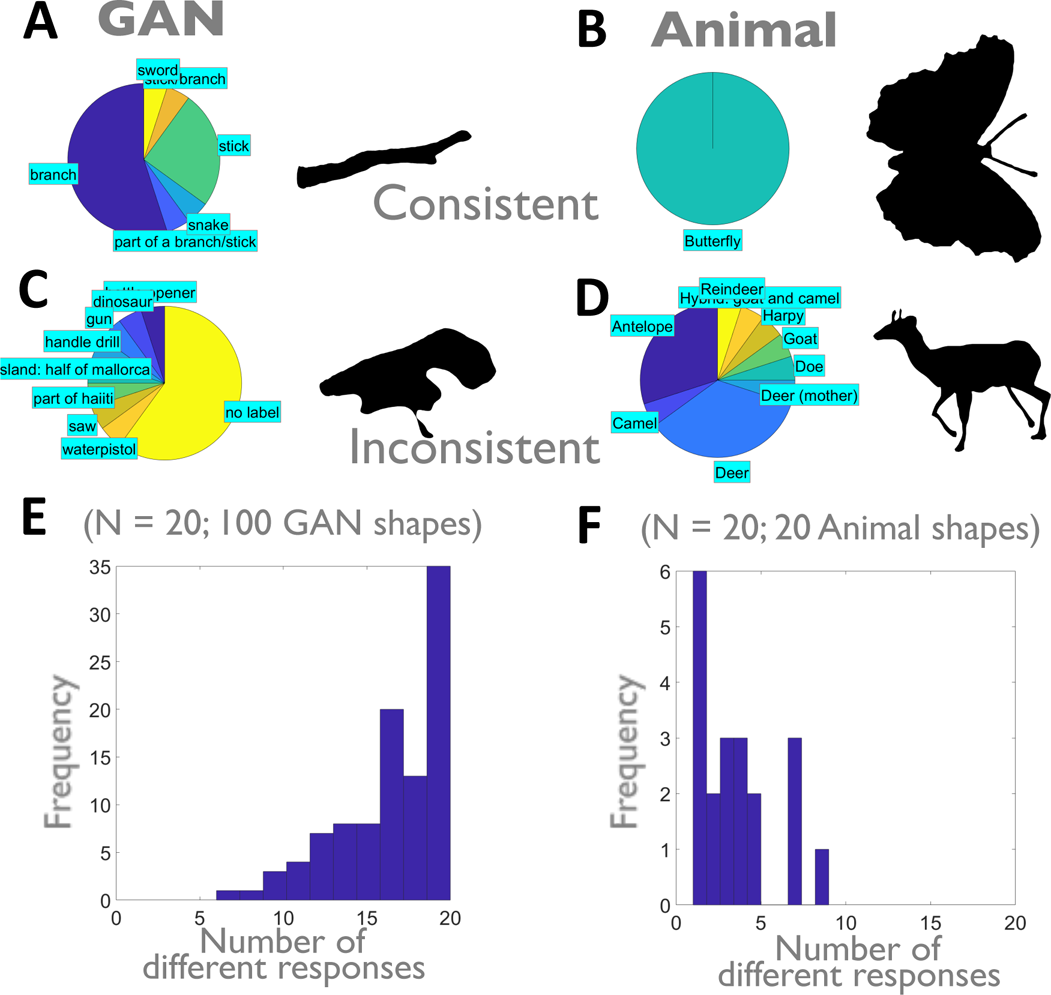
GAN category identification shows human uncertainty in object class. The most consistent (**A,B**) and inconsistent (**C,D**) category responses (N=20) across 100 GAN shapes and 20 animal shapes. Histogram of different responses across **(E)** GAN and **(F)** animal shapes. Human observer responses tended to be different across GAN shapes, and the same across animal shapes. Animal-like category responses to GAN images are much more uncertain than for animal shapes. (See Supplemental Methods for experimental procedure)

**Figure S6.**
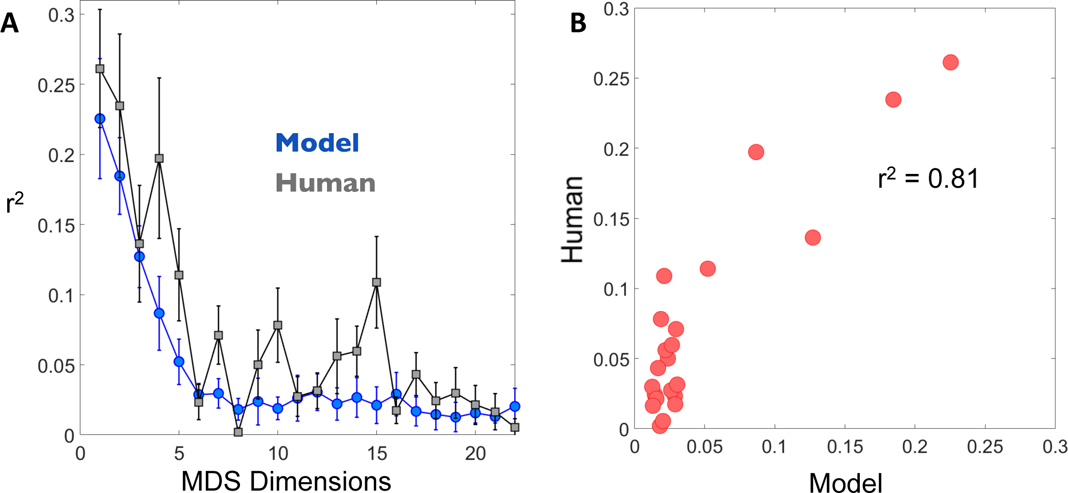
Correlation of human shape similarity ratings with fall-off of variance in MDS dimensions for animal shapes. (A) Variance accounted for in ShapeComp by animal shapes (in blue) and for human similarity ratings (in grey) across pairs of GAN shapes. Error bars for animal data indicate standard deviation across 10 sets of 500 different animals shapes. Error bars for human data indicate standard deviation across observers (N = 14). **(B)** Scatterplot of average human versus model for 22 MDS dimension. ShapeComp and human are highly related.

**Figure S7.**
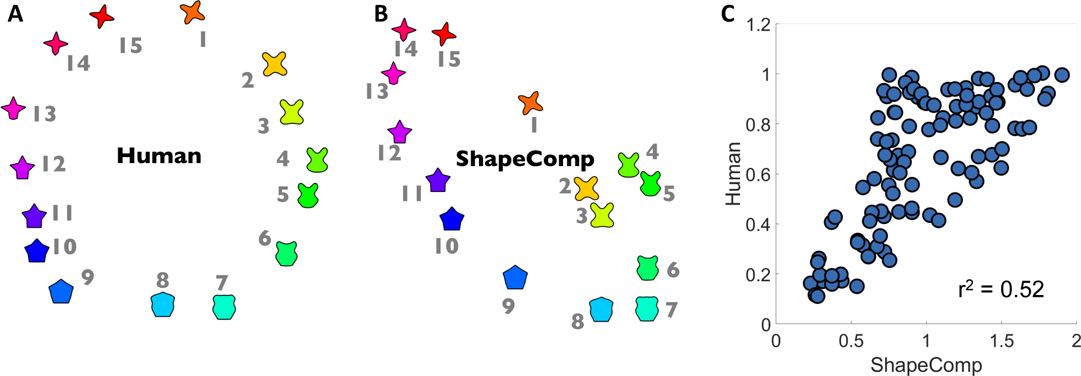
C**o**mparing **ShapeComp and humans across the validated circular shape space set** (from Li et al., [2]). **(A)** Human Data from Li et al. **(B)** Predictions of ShapeComp. Note that while ShapeComp’s arrangement in is more elliptical, there are many similarities with humans. ShapeComp correctly predicts (i) large gaps between shapes 1 and 15, 1 and 2, and 8 and 9, (ii) the circular nature of the data set, (iii) subjective difference between 1 and 11 is smaller than between 14 and 8, yielding the elongated arrangement. **(C)** Correlation between ShapeComp and human similarity judgments for the distances between all possible (105 pairs) (*r*^2^ = 0.5249, *p* <0.01). Given the noise uncertainty across observers – which is unknown for the circular shape set - ShapeComp appears to be a good model of human behaviour. Note, given the symmetry in the circular shape set, here the first point across shapes that were inputted into ShapeComp22 was set to be nearest the same spatial location rather than the smallest x-value.

**Figure S8.**
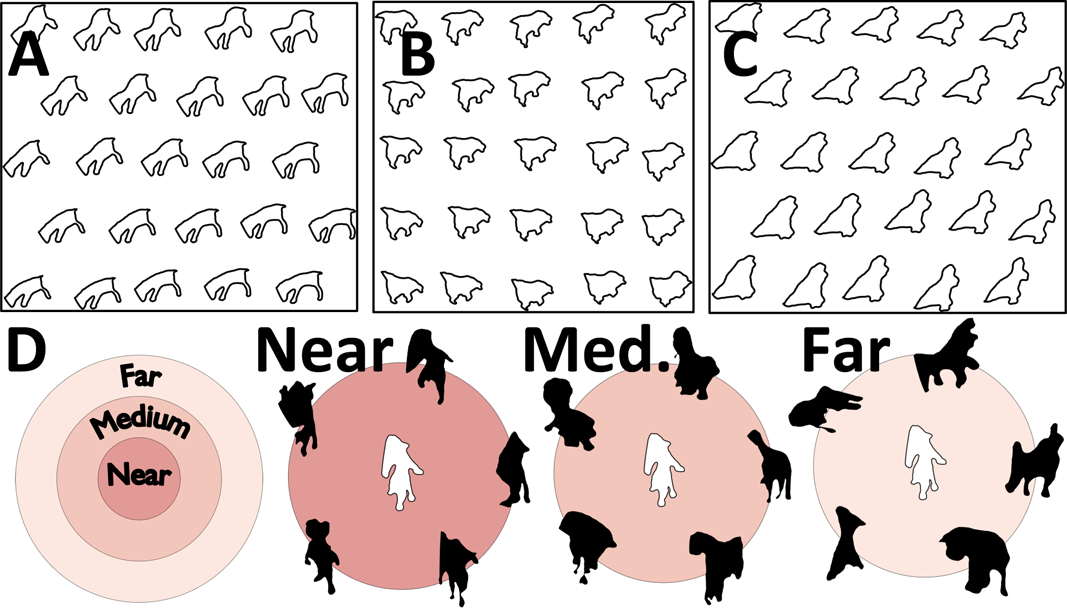
Shape spaces. ShapeComp paired with GAN can be used to create perceptual uniform shape spaces (**A-C**) along a hexagonal **(A, C)** or uniform **(B)** grid or in selecting test shapes that have similar shape similarities **(D**, near, medium, or far in terms of their distances in ShapeComp) to the central sample shape.

**Figure S9.**
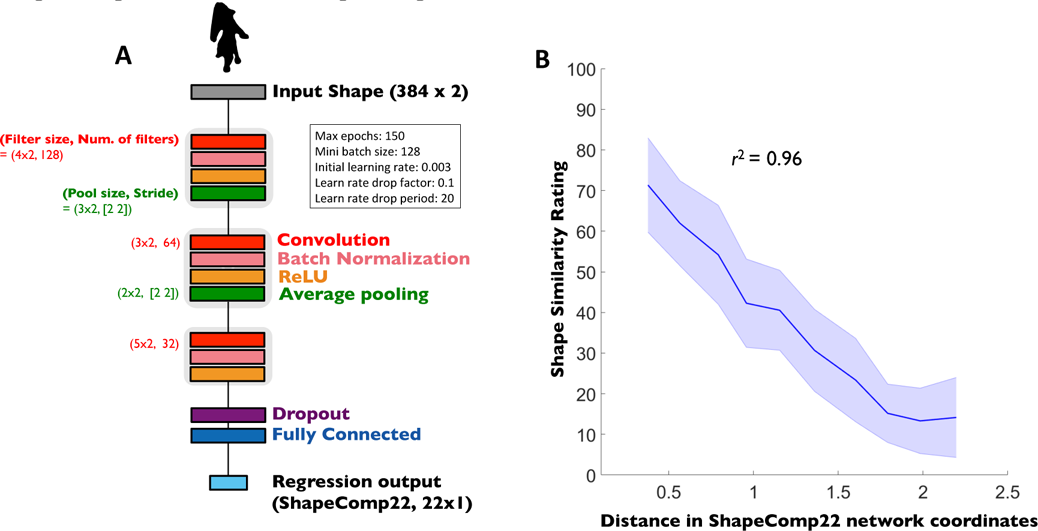
Computing ShapeComp coordinates quickly with a neural network. (A) ShapeComp22 neural network (trained on 800,000 shapes) gets as input a 384×2 shape and outputs the 22D high-dimensional shape space that is **(B)** highly predictive of the human shape similarity ratings for un-trained shapes (re-plotted from Figure 3B), and captures the distance relationship between pairwise combinations of 1000 shapes (*r*^2^ = 0.87, *p* <0.01)

### Supplemental Tables

**Table S1.** **List of 109 shape descriptors in ShapeComp.**

| Descriptors | Additional Comments |
| --- | --- |
| **Simple shape descriptors** | Motivated by Zhang and Lu (2004), Paulun et al. (2015), and Peura and Iivarinen (1997). Many of these descriptors were computed with MATLAB's regionprops command. Computations for (1)-(22) are based on image representation of shape. (23-109) are based on point representation of shape. (27) and (28) are computed with help of geom2d toolbox (by David Legland) from the MATLAB file exchange. |
| (1) Area | regionprops |
| (2) Perimeter | regionprops |
| (3) Eccentricity | regionprops |
| (4) Major Axis Orientation | regionprops |
| (5) Ratio of principle axis | Major/minor axis length |
| (6) Extent | regionprops |
| (7) Compactness | $\frac{area^2}{2\pi\sqrt{MajorAxisLength^2 + MinorAxisLength^2}}$ |
| (8) Solidity |  |
| (9) Circularity A | $4\pi(Area/Perimeter^2)$; from Paulun et al. (2015) based on regionprops output. |
| (10) Circularity B | $Perimeter^2 / Area$ ; from Zhang and Lu (2004) |
| (11) Convexity | Ratio of perimeters of the convex hull over that of the original contour |
| (12) Centroid A | X-coordinate of shape centroid in image (from Paulun et al. (2015)) |
| (13) Centroid B | Y-coordinate of shape centroid in image (from Paulun et al. (2015)) |
| (14) Circularity Ratio | Area of shape to area of circle, where the circle has the same perimeter |
| (15) Convex Area | Paulun et al. (2015) regionprops |
| (16) Area/Perimeter | Zhang and Lu (2004) |
| (17-19) Statistics for horizontal (x) distribution of image intensities | Standard deviation, skewness, kurtosis; from Paulun et al. (2015) |
| (20-22) Statistics for vertical (y) distribution of image intensities | Standard deviation, skewness, kurtosis; from Paulun et al. (2015) |
| (23-26) Curviness at different scales | RMS error of shape to blurred version of itself. Blurred versions were filtered version of original shape with kernel size of 2, 4, 8, or 16 points. Based on Paulun et al. (2015). |
| (27) Circular Variance | Peura and Iivarinen, 1997 |
|  | The proportional mean square error with respect to a circle of the same area |
| (28) Elliptical Variance | Peura and Iivarinen, 1997<br>The proportional mean square error with respect to an ellipse of the same area |
| **Shape Context** | Belongie and Malik (2000) |
| (29) Shape Context | A joint histogram of distance (bins = 15) and angle (bins = 15) |
| (30) Histogram of chord lengths | bins = 100 |
| (31) Histogram of chord angles | bins = 100 |
| **Shape Signatures** | Motivated by Zhang and Lu (2004). |
| (32) Radius Signature |  |
| (33) Curvature Signature |  |
| (34) Triangular Area Signature |  |
| (35) Tangent Angle Signature |  |
| (36) Cumulative Tangent Angle Signature |  |
| (37) Horizontal (x) distribution of points |  |
| (38) Vertical (y) distribution of points |  |
| **Frequency Decomposition** |  |
| (39 - 48) Coefficients 1 to 10 | Separate descriptors for first 10 coefficients; Kuhl and Giardini (1982) |
| (49) Complete Fourier Description | Kuhl and Giardini (1982) |
| **Surprisal** | Information, in Shannon's sense, along contours<br>Feldman and Singh (2005). |
| (50) Signed |  |
| (51) Unsigned |  |
| **Boundary Moments** | Mean, standard deviation, skew, and kurtosis of (30-38). Motivated by Zhang and Lu (2004). |
| (52-55) Statistics for histogram of chord lengths |  |
| (56-59) Statistics for histogram of chord angles |  |
| (60-63) Statistics for radius signature |  |
| (64-67) Statistics for curvature signature |  |
| (68-71) Statistics for triangular area signature |  |
| (72-75) Statistics for tangent angle |  |
| (76-79) Statistics for cumulative tangent angle |  |
| (80-83) Statistics for x-values |  |
| (84-87) Statistics for y-values |  |
| **Shape Skeleton** | Feldman and Singh (2006)<br>Wilder, Feldman, and Singh (2011)<br>Computed with ShapeToolbox |
| (88) Total number of skeletal branches |  |
| (89) Maximum depth of skeleton |  |
| (90) Mean skeletal depth |  |
| (91) mean branch angle |  |
| (92) mean distance along each parent axis at which each child stems |  |
| (93) mean length of each axis relative to root |  |
| (94) total absolute unsigned turning angle integrated along the curve |  |
| (95) total (absolute value of) signed turning angle of each axis integrated along the curve |  |
| (96) standard deviation of (91) | To our knowledge, skeletal summaries (96-102) have been previously used to describe shape. |
| (97) standard deviation of (92) |  |
| (98) standard deviation of (93) |  |
| (99 – 102) Statistics for distribution of ribs | Mean, standard deviation, skewness, kurtosis |
| **Fourier Component** | Fourier descriptions of shape signatures from (32-38) in terms of absolute value of one-sided Fourier component |
| (103) Fourier transform of radius signature |  |
| (104) Fourier transform of curvature signature |  |
| (105) Fourier transform of triangular area signature |  |
| (106) Fourier transform of tangent angle signature |  |
| (107) Fourier transform of cumulative tangent angle |  |
| (108) Fourier transform of distribution of X values |  |
| (109) Fourier transform of distribution of Y values |  |

## References

1. I. Biederman, Recognition-by-components: a theory of human image understanding. Psychological review, 94(2), 115 (1987).

2. D. Marr, Nishihara, H. K, Representation and recognition of the spatial organization of three-dimensional shapes. Proceedings of the Royal Society of London. Series B. Biological Sciences, 200(1140), 269–294, (1978).

3. A. Pentland, Perceptual organization and the representation of natural form. Artif. Intell. 28, 293–331 (1986a).

4. B. Landau, L. B. Smith, S. S. Jones, The importance of shape in early lexical learning. Cognitive Development, 3(3), 299–321 (1988).

5. P. Baingio, K. Deiana. Material properties from contours: New insights on object perception. Vision research 115, 280–301, (2015):.

6. V.C. Paulun, T. Kawabe, S.Y. Nishida, R.W. Fleming, Seeing liquids from static snapshots. Vision research, 115, 163–174, (2015).

7. V.C. Paulun, F. Schmidt, J.J.R. van Assen, R.W. Fleming, Shape, motion, and optical cues to stiffness of elastic objects. Journal of vision, 17(1), 20–20, (2017).

8. J.J.R. van Assen, P. Barla, R.W. Fleming, Visual features in the perception of liquids. Current biology, 28(3), 452–458, (2018).

9. F. Schmidt, The Art of Shaping Materials. Art & Perception, 1(aop), 1–27, (2019). https://doi.org/10.1163/22134913-20191116

10. M. Leyton, Symmetry, causality, mind. MIT Press, (1992).

11. P. Spröte, F. Schmidt, R.W. Fleming, Visual perception of shape altered by inferred causal history. Scientific reports, 6, 36245, (2016).

12. F., Schmidt, R.W. Fleming Visual perception of complex shape-transforming processes. Cognitive Psychology, 90: 48–70, (2016).

13. R.W. Fleming, F. Schmidt, Getting “fumpered”: Classifying objects by what has been done to them. Journal of Vision,19(4):15, 1–12, (2019). DOI: 10.1167/19.4.15

14. O. Eloka, V.H. Franz, Effects of object shape on the visual guidance of action. Vision Research 51(8):925–931 (2011).

15. U. Kleinholdermann, V.H. Franz, K.R. Gegenfurtner KR, Human grasp point selection. Journal of Vision 13(8):23–23 (2013)

16. L.K. Klein, G. Maiello, V.C. Paulun, R.W. Fleming, How humans grasp three-dimensional objects. bioRxiv, 476176. DOI: https://doi.org/10.1101/476176 (2019).

17. R.H. Cuijpers, E. Brenner, J.B.J. Smeets JBJ. Grasping reveals visual misjudgements of shape. Experimental Brain Research 175(1):32–44 (2006).

18. L.F. Schettino, S.V. Adamovich, H. Poizner, Effects of object shape and visual feedback on hand configuration during grasping. Experimental Brain Research 151(2):158–166 (2003).

19. G. T., Toussaint (Ed.). Computational morphology: a computational geometric approach to the analysis of form (Vol. 6). Elsevier (2014).

20. F. Ambellan, H. Lamecker, C. von Tycowicz, S. Zachow. Statistical Shape Models-Understanding and Mastering Variation in Anatomy. To appear in: Advances in Experimental Medicine and Biology - Biomedical Visualisatio (2019).

21. P. G. Mezey, Shape-similarity measures for molecular bodies: A 3D topological approach to quantitative shape-activity relations. Journal of chemical information and computer sciences, 32(6), 650–656 (1992).

22. M. Schmittbuhl, B. Allenbach, J.M. Le Minor, A. Schaaf, Elliptical descriptors: some simplified morphometric parameters for the quantification of complex outlines. Mathematical geology, 35(7), 853–871 (2003).

23. P. Ranta, T.O.M. Blom, J.A.R.I. Niemela, E. Joensuu, M. Siitonen, M, The fragmented Atlantic rain forest of Brazil: size, shape and distribution of forest fragments. Biodiversity & Conservation, 7(3), 385–403. (1998).

24. D. Zhang, G. Lu, Review of shape representation and description techniques. Pattern Recognition, 37, 1–19. (2004).

25. S. Biasotti, A. Cerri, A. Bronstein, M. Bronstein, Recent trends, applications, and perspectives in 3d shape similarity assessment. Computer Graphics Forum, 35(6), 87–119 (2016).

26. J.H. Elder, S. Trithart, G. Pintilie, D. MacLean, Rapid processing of cast and attached shadows. Perception, 33(11), 1319–1338 (2004).

27. J.H. Elder, Shape from contour: Computation and representation. Annual review of vision science, 4, 423–450 (2018).

28. M. Peura, J. Iivarinen, Efficiency of simple shape descriptors, Proceedings of the Third International Workshop on Visual Form, Capri, Italy, May, pp. 443–451 (1997).

29. M.A. Rahman, Y. Wang, Optimizing intersection-over-union in deep neural networks for image segmentation. In International symposium on visual computing (pp. 234-244). Springer, Cham (2016, December).

30. S. Belongie, J. Malik, “Matching with Shape Contexts”. IEEE Workshop on Contentbased Access of Image and Video Libraries (CBAIVL-2000). (2000).

31. H. Asada, M. Brady, The curvature primal sketch. IEEE transactions on pattern analysis and machine intelligence, (1), 2–14, (1986).

32. J. Feldman, M. Singh, Bayesian estimation of the shape skeleton. Proceedings of the National Academy of Sciences, 103(47), 18014–18019, (2006).

33. F.P. Kuhl D.R. Giardina, Elliptic Fourier features of a closed contour. Computer Graphics and Image Processing 18: 236–258. (1982).

34. Acharya, T., & Ray, A. K. (2005). Image processing: principles and applications. John Wiley & Sons.

35. Kriegeskorte, N., & Mur, M. (2012). Inverse MDS: Inferring dissimilarity structure from multiple item arrangements. Frontiers in psychology, 3, 245.

36. Charest, I., Kievit, R. A., Schmitz, T. W., Deca, D., & Kriegeskorte, N.(2014). Unique semantic space in the brain of each beholder predicts perceived similarity. Proceedings of the National Academy of Sciences of the United States of America, 111, 14565–14570. http://dx.doi.org/10.1073/pnas.1402594111

37. Jozwik, K. M., Kriegeskorte, N., & Mur, M. (2016). Visual features as stepping stones toward semantics: Explaining object similarity in IT and perception with non-negative least squares. Neuropsychologia, 83, 201– 226. http://dx.doi.org/10.1016/j.neuropsychologia.2015.10.023

38. Bracci, S., & de Beeck, H. O. (2016). Dissociations and associations between shape and category representations in the two visual pathways. Journal of Neuroscience, 36(2), 432–444.

39. Morgenstern, Y., & Kersten, D. J. (2017). The perceptual dimensions of natural dynamic flow. Journal of vision, 17(12), 7–7.

40. Karimpur, H., Morgenstern, Y., & Fiehler, K. (2019). Facilitation of allocentric coding by virtue of object-semantics. Scientific reports, 9(1), 6263.

41. Frey, B. J., & Dueck, D. (2007). Clustering by passing messages between data points. science, 315(5814), 972–976.

42. Montavon, G., Samek, W., & Müller, K. R. (2018). Methods for interpreting and understanding deep neural networks. Digital Signal Processing, 73, 1–15.

43. Ghorbani, A., Abid, A., & Zou, J. (2019, July). Interpretation of neural networks is fragile. In Proceedings of the AAAI Conference on Artificial Intelligence (Vol. 33, pp. 3681–3688).

44. Shepard, R. N., & Cermak, G. W. (1973). Perceptual-cognitive explorations of a toroidal set of free-form stimuli. Cognitive Psychology, 4(3), 351–377.

45. Cortese, J. M., & Dyre, B. P. (1996). Perceptual similarity of shapes generated from fourier descriptors. Journal of Experimental Psychology: Human Perception and Performance, 22(1), 133.

46. Op de Beeck, H.P. Wagemans, J, Vogels, R. Inferotemporal neurons represent low-dimensional configuration of parametrized shapes. Nature Neuroscience, 4(12), 1244, (2001).

47. Op de Beeck, H. P., Torfs, K., & Wagemans, J. (2008). Perceived shape similarity among unfamiliar objects and the organization of the human object vision pathway. Journal of Neuroscience, 28(40), 10111–10123.

48. Op de Beeck, H P, Wagemans, J, Vogels, R, 2008 “The representation of perceived shape similarity and its role for category learning in monkeys: A modeling study” Vision Research 48 598–610

49. Panis, S., Vangeneugden, J., & Wagemans, J. (2008). Similarity, typicality, and category-level matching of morphed outlines of everyday objects. Perception, 37(12), 1822–1849.

50. Yue, X., Biederman, I., Mangini, M. C., von der Malsburg, C., & Amir, O. (2012). Predicting the psychophysical similarity of faces and non-face complex shapes by image-based measures. Vision research, 55, 41–46.

51. Morgenstern, Y., Schmidt, F., & Fleming, R. W. (2019). One-shot categorization of novel object classes in humans. Vision research, 165, 98–108.

52. Destler, N., Singh, M., & Feldman, J. (2019). Shape discrimination along morph-spaces. Vision research, 158, 189–199.

53. Li, A. Y., Liang, J. C., Lee, A. C., & Barense, M. D. (2019). The validated circular shape space: Quantifying the visual similarity of shape. Journal of Experimental Psychology: General.

54. Wilder, J., Fruend, I., & Elder, J. H. (2018). Frequency tuning of shape perception revealed by classification image analysis. Journal of vision, 18(8), 9–9.

55. Shepard, R. N. (1987). Toward a universal law of generalization for psychological science. Science, 237(4820), 1317–1323.

56. Wilder, J., Feldman, J., & Singh, M. (2011). Superordinate shape classification using natural shape statistics. Cognition, 119, 325–340.

57. F. Cutzu and S. Edelman, Faithful representation of similarities among three-dimensional shapes in human vision, Proceedings of the National Academy of Science 93, pp. 12046{12050, 1996.

58. Clément, M., & Elder, J. H. (2019). What are the sparse components of 2D shapes?. reconstruction, 1, 1.

59. Geisler, W.S., Najemnik, J. & Ing, A.D. (2009) Optimal stimulus encoders for natural tasks. Journal of Vision, 17, 1–16.

60. Burge, J. & Geisler, W.S. (2011) Optimal defocus estimation in individual natural images. Proceedings of the National Academy of Sciences, 108 (40): 16849–16854.

61. Burge J, Geisler WS (2015). Optimal speed estimation in natural image movies predicts human performance. Nature Communications, 6: 7900, 1–11 doi:10.1038/ncomms8900

62. Huang, P., Hilton, A., & Starck, J. (2010). Shape similarity for 3D video sequences of people. International Journal of Computer Vision, 89(2-3), 362–381.

63. Hilaga, M., Shinagawa, Y., Kohmura, T., & Kunii, T. L. (2001, August). Topology matching for fully automatic similarity estimation of 3D shapes. In Proceedings of the 28th annual conference on Computer graphics and interactive techniques (pp. 203-212). ACM.

64. Pasupathy, A., El-Shamayleh, Y. and Popovkina, D. V. (2018). Visual shape and object perception, in: *Oxford Research Encyclopedias*, Oxford University Press, Oxford, UK. doi: 10.1093/acrefore/9780190264086.013.75.

65. Bai, X., Liu, W., & Tu, Z. (2009). Integrating contour and skeleton for shape classification, in: International Conference on Computer Vision Workshops (ICCV Workshops), IEEE, pp. 360–367.

66. Goodfellow I, et al. (2014) Generative Adversarial Nets. Advances in Neural Information Processing Systems 27, pp 2672–2680.

67. Radford A, Metz L, Chintala S (2015) DCGAN: Unsupervised Representation Learning with Deep Convolutional Generative Adversarial Networks. arXiv 1511.06434. https://arxiv.org/pdf/1511.06434.pdf

68. Brainard, D. H. (1997). The psychophysics toolbox. Spatial vision, 10(4), 433–436.

69. Kleiner, M. et al. What’s new in Psychtoolbox-3? Perception 36, S14 (2007).

## Supplemental References

1. Bai, X., Liu, W., & Tu, Z. (2009). Integrating contour and skeleton for shape classification, in: International Conference on Computer Vision Workshops (ICCV Workshops), IEEE, pp. 360–367.

2. Li, A. Y., Liang, J. C., Lee, A. C., & Barense, M. D. (2019). The validated circular shape space: Quantifying the visual similarity of shape. Journal of Experimental Psychology: General.

3. Zhang, D., & Lu G. (2004). Review of shape representation and description techniques. Pattern Recognition, 37, 1–19.

4. V.C. Paulun, T. Kawabe, S.Y. Nishida, R.W. Fleming, Seeing liquids from static snapshots. Vision research, 115, 163–174, (2015).

5. Peura, M. & Iivarinen, J. (1997). Efficiency of simple shape descriptors, in: Proceedings of the Third International Workshop on Visual Form, Capri, Italy, May, pp. 443–451.

6. Belongie, S. & Malik, J. (2000). “Matching with Shape Contexts”. IEEE Workshop on Contentbased Access of Image and Video Libraries (CBAIVL-2000).

7. Kuhl, F. P., & Giardina, D. R. (1982). Elliptic Fourier features of a closed contour. Computer Graphics and Image Processing 18: 236–258.

8. Feldman and Singh (2005). Information along contours and object boundaries. Psychological Review. Vol 122. No. 1, 243–252

9. Feldman, J., & Singh, M. (2006). Bayesian estimation of the shape skeleton. Proceedings of the National Academy of Sciences, 103(47), 18014–18019.

10. Wilder, J., Feldman, J., & Singh, M. (2011). Superordinate shape classification using natural shape statistics. Cognition, 119, 325–340.

